# Molecular Characterization of the Tumor Microenvironment in Renal Medullary Carcinoma

**DOI:** 10.1101/2022.04.28.489873

**Authors:** David S. Tourigny, Mark Zucker, Minsoo Kim, Paul Russo, Jonathan Coleman, Chung-Han Lee, Maria I. Carlo, Ying-Bei Chen, A. Ari Hakimi, Ritesh R. Kotecha, Ed Reznik

**Affiliations:** Irving Institute for Cancer Dynamics, Columbia University, Schermerhorn Hall, 1190 Amsterdam Ave, New York, NY 10027, USA; School of Mathematics, University of Birmingham, Birmingham, B15 2TT, UK; Department of Epidemiology and Biostatistics, Sloan Kettering Institute, Memorial Sloan Kettering Cancer Center, New York, NY; Department of Surgery, Memorial Sloan Kettering Cancer Center, New York, NY; Department of Medicine, Memorial Sloan Kettering Cancer Center, New York, NY; Genitourinary Oncology Service, Department of Medicine, Memorial Sloan Kettering Cancer Center, New York, NY; Department of Pathology, Memorial Sloan Kettering Cancer Center, New York, NY

## Abstract

Renal medullary carcinoma (RMC) is a highly aggressive disease associated with sickle hemoglobinopathies and universal loss of the tumor suppressor gene *SMARCB1*. RMC has a relatively low rate of incidence compared with other renal cell carcinomas (RCCs) that has hitherto made molecular profiling difficult. To probe this rare disease in detail we performed an in-depth characterization of the RMC tumor microenvironment using a combination of genomic, metabolic and single-cell RNA-sequencing experiments on tissue from a representative untreated RMC patient, complemented by retrospective analyses of archival tissue and existing published data. Our study of the tumor identifies a heterogenous population of malignant cell states originating from the thick ascending limb of the Loop of Henle within the renal medulla, displaying the hallmarks of increased resistance to cell death by ferroptosis and proteotoxic stress driven by *MYC*-induced proliferative signals. Specifically, genomic characterization of RMC tumors provides substantiating evidence for the recently proposed dependence of *SMARCB1*-difficient cancers on an intact *CDKN2A*-p53 pathway and we suggest increased cystine-mTORC-*GPX4* signaling also plays a role within transformed RMC cells. We further propose that RMC has an immune landscape comparable to that of untreated RCCs, including heterogenous expression of the immune ligand *CD70* within a sub-population of tumor cells, which could provide an immune-modulatory role that serves as a viable candidate for therapeutic targeting.

## Introduction

Renal medullary carcinoma (RMC) is a rare (less than 0.5% of all renal carcinomas) yet highly aggressive kidney malignancy among adolescents and young adults (median age of incidence of 28 years old (1)(2)), which is almost uniformly associated with sickle cell hemoglobinopathies such as sickle cell trait (SCT) and less commonly sickle cell disease (SCD), or thalassemia (3)(4). Fewer than 5% of patients with RMC survive longer than 36 months and the disease is highly resistant to conventional renal cell carcinoma (RCC) therapies, producing only a brief response in approximately 29% of cases (1)(2). A better understanding of RMC biology is therefore an essential prerequisite for developing effective treatment strategies, but so far progress has been hampered due to a paucity of molecular information about this rare disease (5).

RMC is universally characterized by loss, as determined by immunohistochemistry (IHC), of protein *SMARCB1/INI1* that forms a subunit of the SWI/SNF chromatin remodeling complex, and molecular profiling studies performed to date have revealed the majority of biallelic inactivation of the *SMARCB1* gene occurs either via concurrent hemizygous loss and translocation or by homozygous loss (6)(7)(8)(9)(10)(11)(12)(13). Outside of *SMARCB1* inactivation, there are no recurrent somatic genetic alterations and the mutational load of RMC appears remarkably sparse compared to other RCCs (13), suggesting that loss of *SMARCB1/INI1* protein expression is both a truncal and potent driver of disease. In addition to RMC, *SMARCB1* alterations are found in approximately 20% of all cancers (14)(15), and biallelic inactivation of SMARCB1 is also a universal characterizing feature of kidney malignant rhabdoid tumor (MRT) that similarly has an extremely low rate of mutation and aggressive nature of onset (16). Whether parallels between RMC and kidney MRT extend to the molecular level remains largely unexplored. It has recently been established that bulk transcriptomic profiles of primary RMC tumors are closest to collecting duct carcinoma (CDC) while remaining relatively distinct from kidney MRT, upper tract urothelial carcinoma (UTUC) and other RCC subtypes including clear cell, papillary, and chromophobe RCC (ccRCC, pRCC and chRCC, respectively) (13).

A proposed mechanism for RMC propagation is attributed to a predisposition for red blood cell (RBC) sickling in the renal medulla of SCT/D patients that triggers chronic hypoxia followed by DNA damage and repair-induced loss of *SMARCB1* (17)(5)(18). Indirect support for this hypothesis comes from the observation that *SMARCB1* is localized on a fragile region of chromosome 22 (19)(20). However, the precise cellular origin of RMC still remains unclear. Recent bulk transcriptomic profiling of primary tumors and *in vitro* experiments with RMC-derived cell lines have also implicated the upregulation of pathways associated with *c-MYC*-induced DNA replication stress (13)(21), and the same bulk transcriptomic analyses suggest that RMC expresses a high degree of correlation with collecting duct tissue and an immune profile distinct from ccRCC. ccRCC is closely associated with mutations that lead to stabilization of hypoxia inducible factors (*HIF-1α* and *HIF-2α*), resulting in an oncologic-metabolic shift thought to be an underlying driver for malignancy (22)(23). Therefore, hypoxia might play an important yet multifaceted role across the varied tumor microenvironments (TMEs) of the kidney.

Single-cell transcriptomic analyses complement bulk multi-omic studies by providing information on cell composition and heterogeneity within the TME, which are important factors for understanding the basic biology, treatment strategies and resistance of disease (24)(25). Unlike ccRCC (constituting 70% of all kidney cancers) for which several single-cell transcriptomic data from primary tumors are now available (26)(27)(28)(29)(30)(31), scarcity of cases and difficulties surrounding pre-operative diagnosis of RMC have hitherto restricted molecular characterization of this rare disease to the few bulk genomic and transcriptomic analyses described above. Here, we report genomic, metabolomic and single-cell transcriptomic characterization of the TME from a representative untreated RMC tumor. Our analysis is complemented by retrospective analyses of archival tissue and published data from several cohorts of RCM patients reported previously. Our in-depth molecular characterization of the RMC TME extends current understanding of this rare and devastating disease by consolidating previous findings at the single-cell level and identifying additional malignant, immune and diagnostic signals that may pave the way for more effective treatment strategies.

## Results

### Genomic characterization of RMC

We performed whole exome sequencing (WES) on tumor tissue from the untreated primary nephrectomy of an RMC patient (hereafter referred to as Patient 1) in parallel with a matched normal blood sample to characterize somatic genomic variation within the tumor (Materials and Methods). Contrasted retrospectively against a background of primary tumors from RMC patients previously treated at Memorial Sloan Kettering Cancer Center and for which genomic characterization is available (10), Patient 1 displayed the conventional hallmarks of RMC including SCT diagnosis, right kidney as tissue of origin (approximately 2/3 cases), and loss of *SMARCB1/INI1* expression as revealed by IHC (Figure 1A)(1)(2). Allele-specific copy number and clonal heterogeneity analysis performed using FACETS (32) estimated a tumor purity of 0.18 and loss of heterozygosity (LOH) via deletion of chromosomes 4, 13, 15, and 22, the latter of which includes the *SMARCB1* gene locus (Figure 1B). Figure 1B also shows that FACETS estimated copy-neutral LOH (loss of one parental allele coupled to duplication of the second) within a large region of chromosome 17 and the entirety of chromosome 18. Analysis of somatic nucleotide variants using WES and MSK-IMPACT (Integrated Mutation Profiling of Actionable Cancer Targets) (33) revealed the tumor contained missense mutations in *PRDM1* (residue 352) and *SLX4* (residue 154) that, although potentially oncogenic are not reported within the NCBI ClinVar database, and an indel mutation of *SMARCB1* at the genomic position corresponding to residue 203 (Figure 1C). The indel results in a frameshift mutation that, combined with LOH of chromosome 22, likely explains biallelic inactivation of *SMARCB1* and loss of expression at the protein level. Searching for gene fusion events in RNA transcripts from bulk tumor and normal tissue (Materials and Methods) did not reveal any evidence of further inactivation via translocation for the mutated *SMARCB1* allele. Outside of these events the mutational landscape of the tumor from Patient 1 was relatively sparse compared with other cancer types, as has previously been reported for RMC (13).

**Figure 1.**
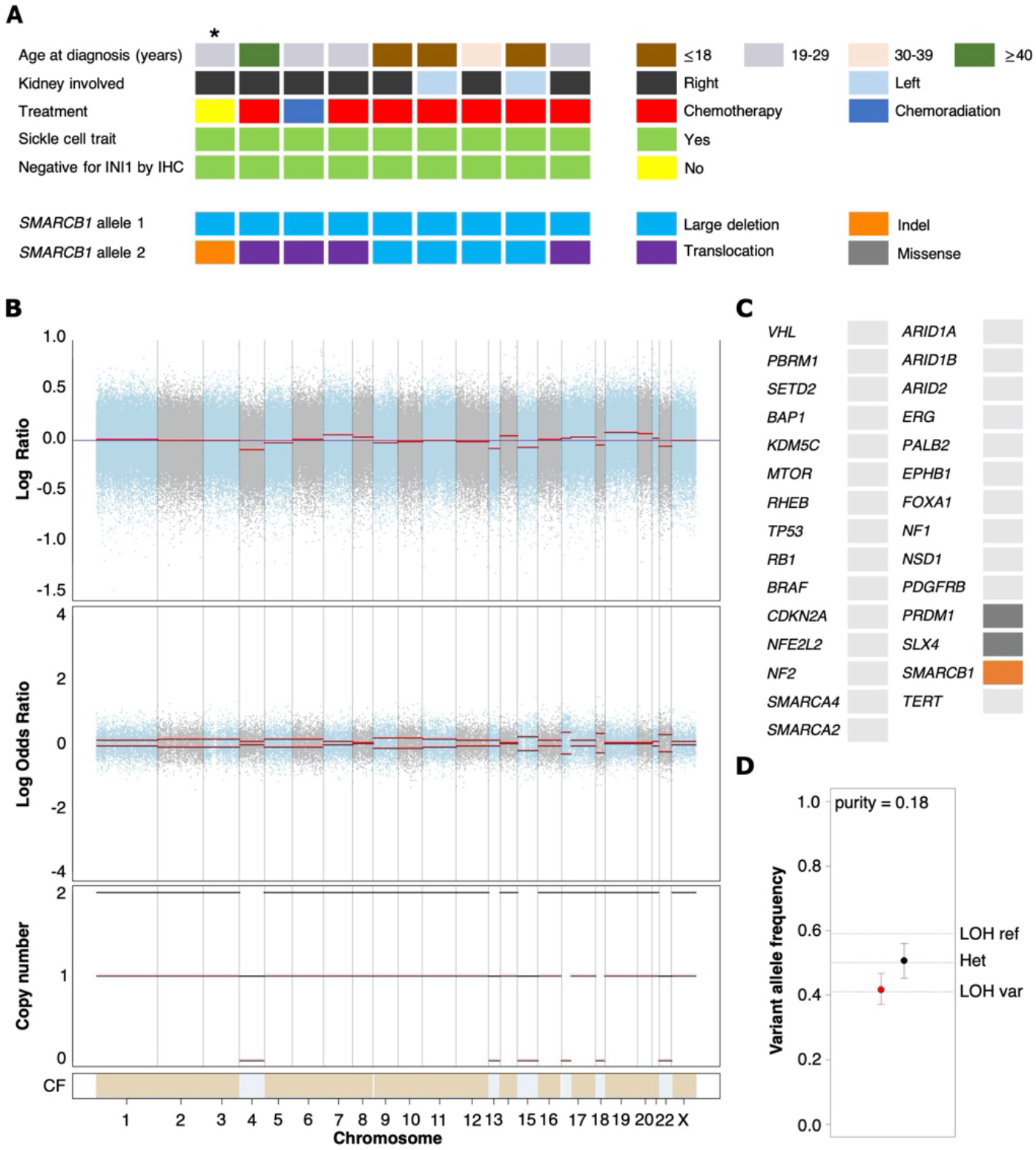
Genomic characterization of primary RMC tumors. (**A**) Oncoplot showing the clinical characteristics and genetic mechanisms of *SMARCB1* inactivation in RMC patients molecularly profiled at Memorial Sloan Kettering Cancer Center. Each column represents a different patient and Patient 1 (first column) from this study is highlighted with an asterisk. (**B**) Integrated visualization of FACETS analysis for WES data from Patient 1. Top two panels display total copy number Log Ratio and allele-specific Log Odds Ratio with chromosomes alternating in blue and grey. Third panel plots the corresponding integer total (black) and minor allele (red) number calls. Estimated cellular fraction (CF) is plotted at bottom in blue with normal diploid state in tan. (**C**) Somatic genomic alterations found in Patient 1 (not including germ line *TP53* allele loss) as reported by MSK-IMPACT analysis, plotted using the same color scheme as in (A). (**D**) Variable allele frequencies (VAFs) of the (likely) pathogenic germ line *TP53* variant inherited by Patient 1 for tumor (red) and normal (black) samples. Vertical bars represent 95% confidence intervals and VAFs corresponding to different allelic configurations of the reference and variant copies of *TP53* are shown as dotted lines. The VAFs of tumor and normal tissue are judged to be significantly different based on a p-value of 0.0189.

Analysis of normal DNA revealed that Patient 1 inherited a G>C germ line variant allele at position 1025 of the reference *TP53* gene, encoding a missense mutation that replaces arginine 342 with a proline residue and consequently results in a physiochemical alteration of p53 at the protein level. This single nucleotide polymorphism (NCBI ClinVar Variation ID 215996) has been annotated as a (likely) pathogenic *TP53* variant segregating with Li Fraumeni syndrome cancers (34) and experimental studies have shown the missense change results in an inactive p53 protein that is defective at both tetramerization and transcriptional transactivation (35). *TP53* is encoded on the region of chromosome 17 that underwent copy-neutral LOH in RMC tissue (Figure 1B), and analysis of the variant allele frequency (VAF) was found to be consistent with loss of the germ line variant allele and duplication of the reference allele within the tumor (Figure 1D). This suggests that the (likely) pathogenic germ line *TP53* variant possibly conferred a selective disadvantage to tumor cells, which implies negative selection against inactivation of *TP53* in RMC. In support of this hypothesis we note that, although the most frequently mutated gene in The Cancer Genome Atlas (TCGA) Pan-Cancer cohort (42% of samples) (36), no somatic *TP53* genetic alteration was observed in any of the RMC tumor samples from Figure 1 nor the 31 RMC patients molecularly profiled in (13). Furthermore, our review of the literature did not reveal any reported case of a primary RMC tumor harboring an oncogenic *TP53* mutation other than Case 10 from (10) (although in that case no detectable structural or copy number alterations involving the *SMARCB1* locus could be detected), and a recent search also previously failed to identify any somatic mutation at the *TP53* locus in MRT (37).

### Heterogenous nature of transformed cell states in RMC

To characterize the cellular composition of the TME in RMC, we performed single-cell RNA sequencing (scRNA-seq) and analysis using the package Seurat (38) on a primary tumor specimen from Patient 1 (Materials and Methods). Our final scRNA-seq dataset included 5610 cells that were parsimoniously annotated by cell type based on the expression of the established cell markers *EPCAM* (epithelial-like cells), *FAP* (fibroblast cells), *CD14* (monocyte/macrophage cells), *CD79A* (B cells), *SDC1* (plasma cells), *CD3D* (T cells), or *CD8A* (CD8+ T cells) (Figure 2A and Supplementary Table 1). We identified proportionally few epithelial-like cells (105 out of 5610), likely reflecting low tumor purity and prominent desmoplasia that is characteristic of RMC (39). Epithelial-like cells further separated into three distinct sub-clusters on the basis of genes identified by differential gene expression analysis (Supplementary Table 2), reflecting their putative role in the malignancy of RMC. Specifically, one sub-cluster (*CXCL14*+ *EPCAM*+ cells) exclusively contained cells expressing the breast and kidney-expressed chemokine *CXCL14* (third top-scoring differentially expressed gene, as ranked by adjusted p-value = 1.566e-9) that is reduced or absent from most cancer cells and used to distinguish normal tissue (40)(41). This finding, together with the observation that the *CXCL14*+ *EPCAM*+ cells retained expression of *SMARCB1* (27.6 % of cells), strongly suggests that the *CXCL14*+ *EPCAM*+ cluster contained either untransformed epithelial cells or those in the early stages of transformation (Figure 2B). When using the top 50 markers as gene sets for cell types identified spatially distributed across the mature human nephron (Figure 2C)(42) to score clusters using the Seurat functions *ModuleScore* and *FindAllMarkers*, the top-ranking (by log fold change) gene set for the *CXCL14*+ *EPCAM*+ cluster corresponded to epithelial progenitor cells (adjusted p-value = 3.621e-17) (Figure 2D).

**Figure 2.**
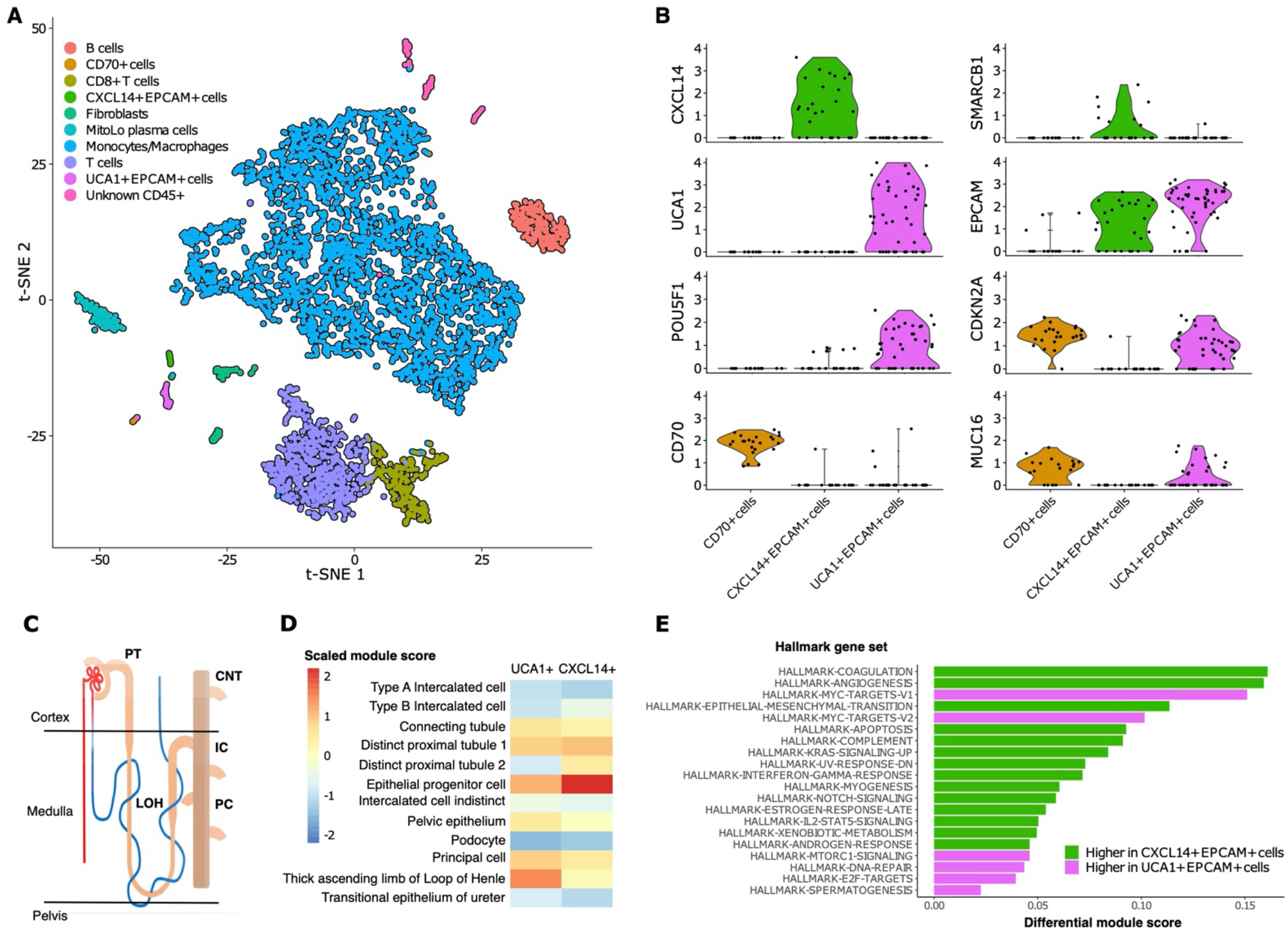
Single-cell composition of RMC. (**A**) t-SNE embedding of transcriptional profiles from all cells (n = 5610). Each dot represents a single cell and colors represent clusters denoted by inferred cell type. (**B**) Normalized expression of genes for the three sub-clusters of epithelial-like cells. Violin plot top and bottom lines indicate range of normalized expression; width indicates number of cells at the indicated expression level. (**C**) Anatomy of the human nephron with site of origin for various cell types. Annotations: PT, proximal tubule; CNT, connecting tubule; LOH, Loop of Henle; IC, intercalated cells; PC, principle cells. (**D**) Module scores (scaled) corresponding to gene set signatures of various epithelial cell types from the mature human kidney for *UCA1+* and *CXCL14+* epithelial-like clusters. (**E**) Differential module scores corresponding to hallmark gene set signatures as calculated between *UCA1+* and *CXCL14+* epithelial-like clusters.

We observed an unequal distribution of *EPCAM* expression among epithelial-like cells identified above: a moderate proportion (62.1%) of cells from the *CXCL14*+ *EPCAM*+ cluster expressed *EPCAM* compared with high (84.6%) and low (12.5%) proportions of cells from the remaining second and third clusters of epithelial-like cells, respectively (Figure 2B). Differential gene expression analysis (Supplementary Table 2) revealed the top-scoring differentially expressed gene of the low *EPCAM* cluster to be *CD70* (adjusted p-value = 1.002e-14)(Figure 2B), which was previously suggested as a target for RCC (43), and also top-scoring differentially expressed genes proposed to be associated with cancer stem cells (e.g. *CD34*, adjusted p-value = 3.163e-14 (44)(45); *CD90/THY1*, adjusted p-value = 3.247e-11 (46)). Conversely, top-scoring differentially expressed genes of the high *EPCAM* cluster contain a large number of ribosomal protein genes, and genes aberrantly expressed in a selection of cancers (e.g., *KRT18*, adjusted p-value = 0.003 (47)(48)) including RCC (e.g., *NAPSA*, adjusted p-value = 4.038e-5 (49)) and RMC (*POU5F1* also known as *OCT3/4*, adjusted p-value = 0.028 (50)) (Figure 2B and Supplementary Table 2). The long non-coding RNA (lncRNA) *urothelial cancer associated 1* (*UCA1*) was also identified among the top-scoring differentially expressed genes of the high EPCAM cluster (adjusted p-value = 3.062e-6) (Figure 2B), substantiating previous findings that this lncRNA shows dramatic upregulation in bulk RMC transcriptome profiles compared to those from adjacent normal tissue (13). Taken in context, this suggests that the *UCA1*+ *EPCAM*+ cluster contained transformed epithelial-like cells further along the RMC oncogenic program. Using the same methodology described above, the top-ranking gene signature for the *UCA1*+ *EPCAM*+ cluster corresponded to the thick ascending limb (TAL) of the Loop of Henle (adjusted p-value = 6.348e-33) (Figure 2D), implicating the TAL as a putative site-of-origin for RMC.

We next customized a version of hallmark gene set analysis (Materials and Methods) to compare differential pathway expression between *CXCL14*+ *EPCAM*+ and *UCA1*+ *EPCAM*+ clusters representing different stages of transformed populations in RMC (Figure 2E). We contrasted single-cell gene set analysis results with the previously reported differential pathway analysis for bulk RMC tumor and adjacent normal tissue (13). Reinforcing the finding that, in bulk, DNA replication stress is a hallmark of RMC, we discovered that the top-five-scoring MSigDB hallmark gene sets representing pathways enriched in the *UCA1*+ *EPCAM*+ cluster to include both sets of *c-MYC* signaling targets (adjusted p-values = 3.096e-6 and 6.434e-5 for target sets *V1* and *V2*, respectively), DNA repair (adjusted p-value = 5.519e-3) and E2F targets (adjusted p-value = 1.990e-2) (Figure 2E). Notably, these results provide new evidence that, at the single-cell level, increased *c-MYC* activity and DNA replication stress can be mapped to transformed epithelial-like cells within the RMC TME. Figure 2E also shows that the mTORC signaling pathway (adjusted p-value = 1.778e-2) was among the top-five-scoring hallmark gene sets upregulated in the *UCA1*+ *EPCAM*+ cells. Taken together, these findings are indicative of increased proliferative and biosynthetic activity following loss of *SMARCB1* within transformed RMC cells.

Epithelial-like cells from the *CD70+* cluster with low *EPCAM* expression levels were enriched for gene set signatures of the interferon gamma and alpha (adjusted p-values = 2.480e-6 and 7.140e-6, respectively), hypoxia (adjusted p-value = 1.532e-4), and inflammatory (adjusted p-value = 2.263e-3) response pathways relative to cells from *UCA1*+ *EPCAM*+ and *CXCL14*+ *EPCAM*+ epithelial-like sub-clusters. To better understand the nature of cells from the *CD70*+ cell cluster, we calculated overall correlation levels between their averaged gene expression profiles with those from the *UCA1*+ *EPCAM+* and *CXCL14*+ *EPCAM*+ sub-clusters (Pearson’s correlation coefficients r = 0.888 and r = 0.831, respectively), revealing that *CD70*+ cells are more closely related to the former even though they have lost many epithelial characteristics, perhaps as a consequence of advanced or divergent progression along the RMC oncogenic program. This claim is further substantiated by the observation that, among all cells, both *UCA1*+ *EPCAM+* and *CD70*+ epithelial-like cells exclusively express *MUC16* (adjusted p-values = 3.883e-132 and 9.967e-295, respectively) (Figure 2B and Supplementary Table 1) as a marker (also known as cancer antigen CA-125) associated with poor prognosis and advanced tumor stage in RCC (51)(52)(53). In summary, our data indicate a heterogenous mixture of at least two or three distinct transformed cell populations within the TME, potentially capturing various stages of the RMC oncogenic program.

### Signals of cystine-dependent ferroptosis resistance in RMC

RBC sickling within the renal medulla is believed to promote chronic hypoxia in RMC (17)(5)(18). Hypoxia and/or ischemia is also thought to increase the susceptibility to cell death via the ferroptosis pathway within transformed epithelial cells, which must therefore ramp up protective mechanisms for their survival (54)(55)(56). In our scRNA-seq analysis we found that among the genes strongly overexpressed in *UCA1+ EPCAM+* versus *CXCL14*+ *EPCAM*+ epithelial-like RMC cell clusters were also those encoding the ferritin light chain (*FTL*, p-value = 8.987e-12), glutathione peroxidase 4 (*GPX4*, p-value = 7.931e-10), and nuclear protein 1 (*NUPR1*, p-value = 0.012), which each play a role in resistance to ferroptosis-induced cell death by sequestering intracellular iron, inhibiting the formation of lipid peroxidases, and transcriptional regulation of iron metabolism, respectively (57)(58)(59).

Overexpression of *FTL, GPX4* and *NUPR1* in *UCA1+ EPCAM+* cells suggests these key genes may be upregulated further along the RMC oncogenic program to increase protection of the tumor against cell death induced by ferroptosis. Correspondingly, the mTORC signaling pathway, which is also upregulated in *UCA1+ EPCAM+* cells (Figure 2D), has recently been shown to couple cystine availability with *GPX4* protein synthesis and regulation of ferroptosis (60). The mTORC signaling mechanism responsible for increasing *GPX4* protein synthesis is dependent on *SLC7A11*-mediated cystine uptake from the extracellular environment, and in bulk RNA-seq data from (13) we found that the cystine transporter *SLC7A11* is expressed at significantly higher levels in RMC tumor versus normal tissue (log2 fold change = 5.267, p-value = 5.386e-14). Although only a handful of epithelial-like RMC cells contain detectable levels of *SLC7A11* transcripts the single-cell level, we also noted that these cells are found exclusively within the *UCA1+ EPCAM+* cluster (data not shown). Taken together, these observations suggest that key components of the newly identified cystine-mTORC-*GPX4* signaling cascade may be mobilized at later stages of the RMC oncogenic transformation program to increase the tumor’s resistance to ferroptosis. This hypothesis would imply that RMC cells are dependent on the uptake of cystine from the extracellular tumor environment.

To probe the metabolic composition of the TME in RMC, we generated bulk metabolomic data from tumor and adjacent normal renal medullary tissue from Patient 1. Due to the absence of statistical power in such a single case study, we analyzed bulk RMC metabolomic data against a background of 140 ccRCC bulk metabolomic data sets (with 71 additional samples from adjacent normal renal cortical tissue) generated and processed using identical protocols (Materials and Methods). To identify individual metabolites found at differential levels of abundance in RMC compared to ccRCC we used the Crawford-Howell test (61) that revealed just two compounds: cystine (Metabolon ID M00056; adjusted p-value = 0.037) and S-Methylcysteine (MeCys) (Metabolon ID M39592; adjusted p-value = 0.038), are found at significantly higher and lower abundance levels, respectively, in the TME of RMC compared to that of ccRCC (Figure 3A and Supplementary Table 3). The same analysis performed on normal samples shows that MeCys, an unconventional amino acid excreted in urine (62), is also found at higher abundance levels in tumor adjacent normal tissue from the renal medulla compared to renal cortex (adjusted p-value = 0.010) (Figure 3B and Supplementary Table 4). Parsimonious interpretation of these results implies that the only significant difference in metabolite abundances attributable to tumor type and not differences in renal site of origin are the significantly lower abundance levels of cystine found in RMC compared to ccRCC. This result could be explained by a relatively higher rate of cystine uptake in RMC, possibly required to support cystine-dependent resistance to ferroptosis.

**Figure 3.**
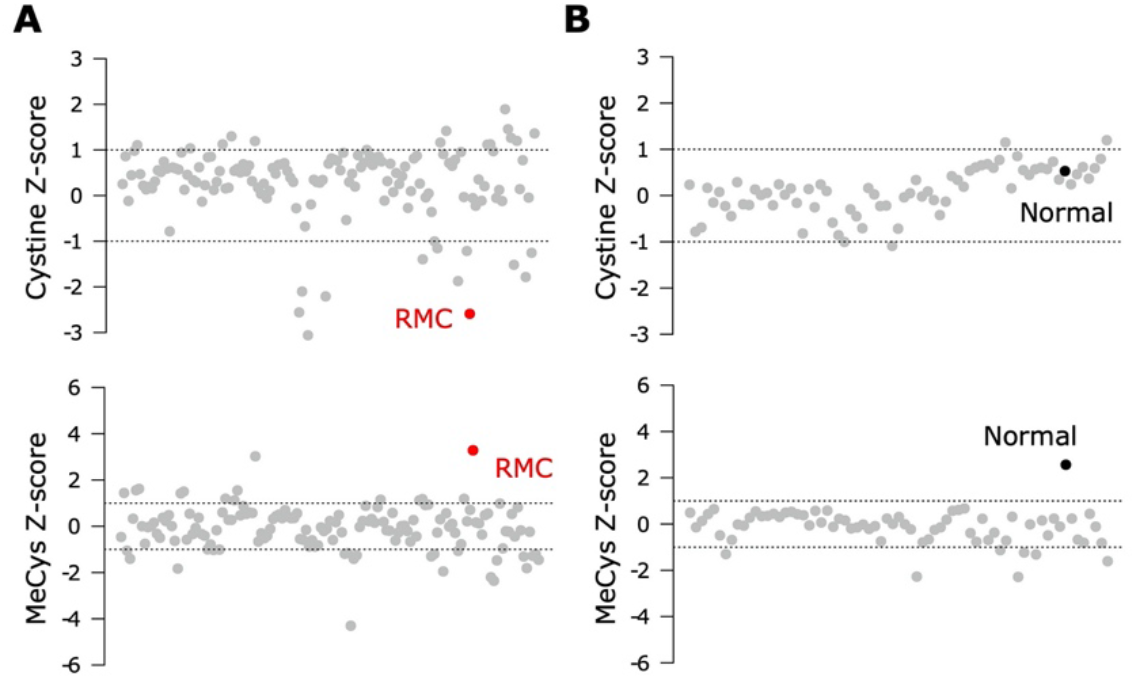
Signals of increased cystine uptake in the RMC primary tumor. (**A**) Z-scores for cystine and MeCys from RMC tumor tissue sample (red) on a background of ccRCC tumor samples (grey). (**B**) Z-scores for cystine and MeCys from normal tissue sample (black) adjacent to an RMC tumor on a background normal tissue (grey) adjacent to ccRCC tumor samples. Dotted lines represent one standard deviation away from the mean.

### Immune composition and crosstalk within the RMC microenvironment

RCCs (and ccRCC in particular) are known to be among the most immune and vascular infiltrated cancer types (63) and can consequently be responsive to various forms of advanced immunotherapy (64). RMC has recently been proposed to have a distinct immune profile compared to other RCCs, based on using bulk RNA-seq deconvolution methods to show that RMC includes an abundance of myeloid and B linage cells compared to ccRCC while retaining a comparably high level of infiltrating T cells and cytotoxic lymphocytes (13). Our scRNA-seq analysis of RMC revealed that the TME was dominated by immune cells from the monocytic lineage (Figure 2A) alongside a complete absence of transcripts for the immune checkpoint ligands PD-L1/2 (programmed cell death ligands) and CD80/86 (cytotoxic T-lymphocyte-associated protein 4 ligands) within transformed epithelial-like cells.

Leveraging the fact that multiple single-cell datasets characterizing the immune landscape of ccRCC across various disease stages and with therapy are now publicly available (29)(30)(31), we developed a correlation-based method to further compare immune cell cluster compositions from RMC to those from ccRCC (Materials and Methods). Using a ccRCC scRNA-seq data set from (31), we evaluated the correlation between average gene expression profiles for each annotated *CD45*-postive cluster from that study with those for each *CD45*-postive cluster from our single-cell RMC dataset (Figure 4A). We found relatively consistent levels of correlation between average immune cell gene expression profiles across clusters of lymphoid lineages from both RCC types, while the RMC Monocyte/Macrophage cluster displayed a graded correlation signature with cell types from the myeloid linages detected in ccRCC. Specifically, Figure 4A shows that the RMC Monocyte/Macrophage cell cluster was most enriched for a transcriptional profile associated with tumor associated macrophages (TAMs) expressing high levels of *GPNMB* that was shown to be associated with the anti-inflammatory M2 macrophage polarization signature found predominantly in untreated ccRCC (31). Concordantly, lower levels of correlation between the RMC Monocyte/Macrophage cell cluster and remaining ccRCC TAM clusters is consistent with the shift towards the pro-inflammatory M1 macrophage polarization signature in treated tumors (29)(30)(31). It remains unclear whether a shift to pro-inflammatory states would similarly be observed within TAM populations from RMC tumors treated using immunotherapy targeting the classical PD-L1/2 or CD80/86 immune checkpoint regulators, which appear relatively absent in this cancer type.

**Figure 4.**
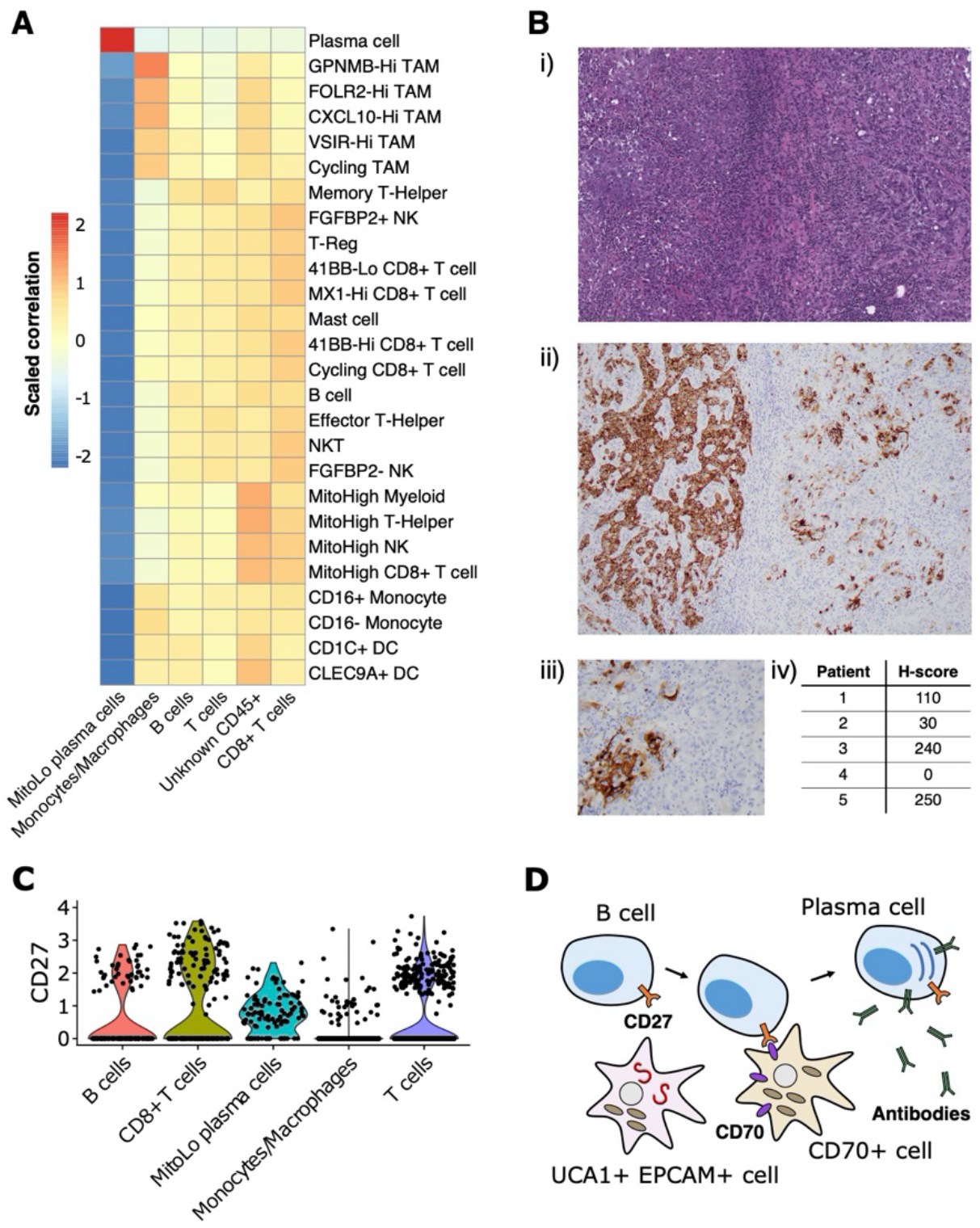
Immune landscape and cross-talk in the RMC tumor microenvironment. (**A**) Correlations between transcriptional profiles of immune cell types identified in RMC and ccRCC, respectively. (**B**) *CD70* immunohistochemistry: (i) representative HE images of tumor regions from Patient 1; (ii) corresponding *CD70* immunostaining of tumor regions from Patient 1; (iii) heterogenous *CD70* immunoreactivity of tumor cells at high magnification; (iv) H-score for five RMC tumor samples (including Patient 1; H-score = 110) assayed for *CD70* protein expression. (**C**) Violin plot displaying normalized expression of *CD27* across immune cell clusters in RMC. (**D**) Cartoon schematic illustrating a putative role for the *CD70*-expressing tumor cell sub-population in co-stimulation of plasma cell differentiation via the *CD27* receptor.

*CD70* has recently been proposed as an emerging target in cancer immunotherapy (65), particularly in the case of RCC where its overexpression in ccRCC is thought to be driven by stabilization of hypoxia inducible factors (43)(66). Using bulk RNA-seq data from (13), we observed that *CD70* mRNA is highly overexpressed in RMC tumor compared with adjacent normal tissue (log2 fold change = 7.724, p-value = 3.269e-23). We rationalized that increased *CD70* expression in RMC could be explained by the presence of *CD70*+ epithelial-like cells or an increase in tumor infiltrating B cells, since *CD70* was identified as a top-scoring differentially expressed gene for both clusters in our scRNA-seq analysis (Supplementary Table 1). Indeed, custom-built gene sets generated using the top 50 scoring differentially expressed genes from both B cell and *CD70*+ cell clusters found in our scRNA-seq data demonstrated comparable gene set enrichment analysis (GSEA) scores in bulk RMC tumor (normalized enrichment score (NES) = 2.382, p-value = 3.461e-8 and NES = 2.002, p-value = 2.798e-5 for gene sets of B and *CD70*+ cells, respectively), suggesting both types of *CD70*-expressing cells are overrepresented in malignant tissue.

To further explore the relevance of *CD70* expression in RMC, we performed IHC staining on primary RMC tumor tissue sections from Patient 1 and four other RMC patients previously treated at MSKCC. Heterogenous intra-tumor *CD70* protein expression on carcinoma (or RMC tumor) cells was found in 80% (4/5; including Patient 1) of the primary tumors tested (as quantified by H-score, Materials and Methods) (Figure 4B), substantiating the discovery that *CD70* is expressed in a sub-population of transformed RMC cells. *CD70* is the only known ligand for the immune receptor *CD27*, and it is therefore conceivable that a sub-population of transformed RMC cells expressing *CD70* performs an immuno-modulatory role by co-stimulation of lymphocytes via the *CD27* receptor. Concordantly, our scRNA-seq data reveal that *CD27* is expressed at moderate levels within T cells (22.65 % of cells), *CD8+* T cells (32.30 % of cells), B cells (17.28 % of cells), and substantially higher levels within the MitoLo plasma cells (60.45 % of cells) (Figure 4C), suggesting significant levels of *CD27*-*CD70*-mediated cross-talk are possible between different cell populations within the RMC TME. In addition to an abnormally low level of mitochondria-encoded transcripts found within the MitoLo plasma cell population (Materials and Methods), the top-scoring hallmark gene set for this cluster corresponds to the unfolded protein response (UPR) pathway (adjusted p-value = 7.320e-29), indicative of a stimulated cell state associated with a dramatic increase in antibody synthesis.

## Discussion

RMC is a rare yet devastating disease uniquely associated with sickle hemoglobinopathies and *SMARCB1* inactivation, whose mechanistic origin still remains largely unexplained because a relative shortage of cases has made molecular profiling difficult. In light of this, our study has presented an in-depth molecular characterization of the TME from an untreated RMC patient carrying a (likely) pathogenic *TP53* germ line allele that was somatically lost in the tumor. Other than this unique case of germ line mutant allele LOH, the genomic landscape of the RMC tumor was generally representative of disease, including biallelic inactivation of *SMARCB1* with a low mutational burden otherwise.

We have mapped transcriptional signals from differentially expressed genes that are characteristic of RMC (13)(21) to distinct sub-populations of transformed renal epithelial cells originating from the TAL. We therefore tentatively put forward the TAL as a putative site-of-origin for RMC. Our single-cell analyses suggest that the transformed RMC cell state expressing lncRNA *UCA1* is associated with increased *c-MYC* activity, DNA replication stress and mTORC signaling compared to untransformed epithelial cells. Although *UCA1* was originally believed to be a highly-specific marker for urothelial cancer (67), subsequent studies revealed its overexpression in a variety of human malignancies where it is understood to promote proliferation, migration and immune escape (68). Our findings provide additional evidence that *UCA1* could serve as a marker gene for RMC malignancy as was previously suggested based on bulk RMC transcriptome data (13). Likewise, bulk transcriptome data and in-vitro cell line experiments also support the increase in *c-MYC* activity and DNA repair stress following *SMARCB1* inactivation in both RMC (13)(21) and MRT (37). The authors of the latter study concluded that a genetically-intact *CDKN2A*-p53 pathway is necessary for regulating the UPR pathway to buffer transformed cells against dramatic elevation of proteotoxic stress that results from activation of *c-MYC* (37).

Our observation that a (likely) pathogenic *TP53* germ line allele was somatically lost in the RMC tumor could be explained by assuming a functional *CDKN2A*-p53 signaling axis protects against cellular stress associated with *SMARCB1* inactivation, as proposed in a previous study (37). In this scenario, high levels of ER stress and proteotoxicity levels requiring modulation by the *CDKN2A*-p53 signaling may have provided a selective advantage to tumor cells that had lost the (likely) pathogenic *TP53* allele. Concordantly, the previous study (37) failed to identify any somatic mutation at the *TP53* locus in MRT and we were likewise unable to find any instance of somatic *TP53* inactivation in RMC tumors that were confirmed *SMARCB1*-defficient at the genomic level. It is therefore possible that engagement of the *CDKN2A*-p53 pathway confers a survival benefit to *SMARCB1*-defficient RMC tumors following elevated *c-MYC* signaling, as previously suggested for MRT (37). We note that this proposal could also explain why *CDKN2A* is only expressed in epithelial-like cell clusters lacking *SMARCB1* transcripts in RMC (Figure 2B). We furthermore presented evidence that resistance to ferroptosis--mediated by the cystine-mTORC-*GPX4* signaling pathway (60)--is an additional trait of the transformed RMC cell state, which could be explained physiologically given the relevance of tissue ischemia and hypoxia for disease. Whether the *CDKN2A*-p53 and cystine-mTORC-*GPX4* signaling axes complement or antagonize one another remains to be explored (69)(70).

In probing the immune landscape of RMC, we demonstrated that heterogenous expression of *CD70* by a distinct sub-population of malignant cells appears to be a general feature of both RMC and ccRCC (43)(66), and could therefore be considered as a viable target for RCC therapy (65). *CD70* is the only known ligand for *CD27*, a tumor necrosis factor (TNF) receptor expressed on immune cells from the lymphoid lineage (T, B and Natural Killer cells) whose stimulation mediates T and B cell activation (71)(72)(73), and is also thought to induce differentiation of the latter into plasma cells (74). We observed a unique population of plasma cells within our single-cell RMC dataset, expressing high levels of *CD27* and hallmarks of increased cellular stress. It is therefore possible that sub-populations of RMC tumor cells expressing *CD70* trigger hyperstimulation of B cells into a tumor-promoting, imbalanced antibody-producing plasma cell state, and in that way evade an effective immune response (Figure 4D). In particular, this would explain why high levels of *CD70* expression (66) as well as immunoglobulin production and active isotype switching has paradoxically been associated with poor prognosis in RCC (75) and, strikingly, why the top-three-scoring differentially expressed genes *IGHG1, IGLV3-1* and *IGHG3* for the MitoLow plasma cell cluster (Supplementary Table 1) serve as a strong negative prognostic marker for RCC (76).

In summary, our work has provided a detailed molecular characterization of the TME from a primary RMC tumor, which complements the few existing molecular profiling resources available for this rare disease. Limitations of the study include the recognition that scRNA-seq and metabolomic data were derived from just a single patient, and that the relatively low tumor purity of RMC could have masked some of the additional transcriptional signals not detectible in single-cell data. Whilst the problem of limited sample size applies generally to single-cell experiments involving primary tumor tissue, they are particularly acute for RMC given the extremely rare occurrence of disease and frequent absence of a pre-operative diagnosis, which makes prior planning of tissue acquisition and experimentation incredibly difficult. To substantiate the analysis of a single-patient study we were careful to validate our results using retrospective analyses of archival tissue and existing published data where available. Further work is required to increase data coverage and firmly establish our initial findings, with an ultimate goal of driving advancements in diagnosis and treatment of RMC.

## Materials and Methods

### Experimental model and subject details

All research activities were pre-approved by the Institutional Ethics Review Board (IRB) at Memorial Sloan Kettering Cancer Center and individuals were required to provide written informed consent to participate in molecular profiling studies. Patient 1 was a 23-year-old female with sickle cell trait and a right kidney tumor who underwent radical nephrectomy prior to treatment and was found to have stage III (pT3N1) disease. Their RMC diagnosis was confirmed by expert genitourinary pathologists (including Y.B.C.). The patient experienced rapid progression of disease and succumbed to disease 14.5 months following surgery. Archival tissue and DNA sequencing data from previous patients with a diagnosis of RMC between 1996 and 2017 at Memorial Sloan Kettering Cancer Center were retrospectively identified from the institutional databases and correspond to those included in a previous study (10).

### Sample collection, tissue dissociation and single-cell suspension

Samples were directly obtained from the operating room during nephrectomy. At the time of specimen extraction, samples of around 1-1.5cm were obtained by the treating surgeon (A. A. H.) from the tumor region and adjacent normal tissue (at least 2cm away from the tumor). Tissue samples were placed in separate containers containing Roswell Park Memorial Institute (RPMI) medium and transported in regular ice to the laboratory with overall transit time less than 1hr from specimen extraction to cell dissociation. Small tissue aliquots (approx. 3mm^3^/each) were separated from samples for bulk tissue profiling, one of each was snap frozen with liquid nitrogen and stored at -80°C for posterior bulk sequencing, while a second aliquot placed in 10% formalin. The remainder of the tumor tissue was kept fresh RPMI medium and dissociated into a cell suspension by first mincing into small pieces and spun (400 g x 10 min) to obtain a pellet. The pellet was firmly dislodged with tapping and incubated with tumor dissociation cocktail: Liberase (Roche): 250 μg/mL, DNAase (Roche): 100 Units/mL (in HBSS), at 37°C for 25-45 min, until good digestion was visually apparent, after which cells were collected and passed through a 100 μm filter. Filtered cells in enzymatic suspension were diluted with cold incomplete RPMI media and spun (400 g x 10 min), repeating this procedure twice to completely remove the tumor dissociation cocktail. The single-cell suspension was frozen in CTL-Cryo ABC media kit until live cell sorting with flow cytometry.

### Paraffin embedding, hematoxylin-eosin (HE) staining, immunohistochemistry and histologic assessment

Samples previously placed in 10% formalin were left in this solution at room temperature for 24 hr, and then washed with phosphate-buffered saline (PBS) solution and placed in 70% ethanol at 4°C for dehydration. Following dehydration, paraffin-embedding was performed and tissue blocks precured. Formaldehyde-fixed paraffin-embedded (FFEP) blocks were sliced and 5 μm-thick slides obtained for staining. Hematoxylin-eosin (HE) staining was used for pathologic review, and morphologic features recorded. These included growth patterns (reticular/yolk sac-like and cribriform, tubulopapillary, infiltrating tubles/cords/individual cells, and solid sheets), stromal changes, rhabdoid cytology, inflammatory infiltrates, and the presence or absence of drepanocytes, necrosis, and mucin. As reticular or yolk sac tumor-like growth and cribriform pattern often overlap, these were combined as one architectural pattern group. Immunohistochemistry (IHC) was performed on FFEP blocks using mouse monoclonal antibodies *SMARCB1/INI1* (Clone 25/BAF47, dilution 1:200, BD Bioscience) and *CD70* (Clone 301731, R&D Systems #MAB2738). *SMARCB1/INI1* staining was scored as retained or lost when compared to internal positive control cells (endothelial/stromal cells and lymphocytes). For *CD70* H-score assessment, fields were chosen at random at x 400 magnification and the staining intensity in malignant cells was scored as 0, 1, 2, or 3 corresponding to the presence of negative, weak, intermediate or strong staining, respectively. The total number of cells in each field and the number of cells stained at each intensity were counted and H-scores calculated using the formula H-score = (% of cells stained at intensity category 1 × 1) + (% of cells stained at intensity category 2 × 2) + (% of cells stained at intensity category 3 × 3). *CD70* IHC was simultaneously performed on archival tumor tissue from Patient 1 and four additional RMC patients reported in (10) (making five samples in total).

### Targeted DNA sequencing analysis

DNA was extracted from the macro-dissected tumor and matched tissue/blood normal samples using QIAmp DNA FFPE Tissue Kit or EZ1 Advanced XL system (Qiagen) according to the manufacturer’s instructions. Tumor and normal DNA was subject to whole exome sequencing (WES) and MSK-IMPACT (33), a deep sequencing assay for cancer-related genes to identify single nucleotide polymorphisms (SNPs) across the genome. Somatic mutations in tumor DNA were called after private germ line variants detected in the paired normal sample were appropriately detected and filtered out. Copy number analysis and tumor purity estimation was conducted using FACETS (32). Variant allele frequencies (VAFs) were calculated by pooling aligned reads from WES and MSK-IMPACT experiments, and differences between VAFs of tumor and normal samples detected with a p-values and confidence intervals based on a proportionally test assuming an underlying binomial distribution of read counts based on allele frequencies.

### RNA extraction, purification and sequencing

Samples snap frozen and stored at -80°C were thawed in regular ice and pretreated using trizol, chlorophorm and cold centrifugation. Total RNA was extracted using the RNeasy mini extraction kit (QIAgen #74101), according to the manufacturer’s instructions. Bulk whole transcriptome RNA-sequencing libraries were prepared with TruSeq Stranded Total RNA library preparation kit (Illumina) and sequenced on a NovaSeq 6000 S2 or S4 flow cell with a sequencing depth of approximately 200-500 million reads per sample.

### Processing and analysis of bulk RNA-seq data

Bulk RNA-seq data generated in this study along with those accessed from (13) were transformed to gene expression count matrices following the protocol outlined in (77). Briefly, RNA-seq raw read sequences were aligned against the human genome assembly hg19 using the HISAT2 software (version 2.2.1) (78). Aligned reads were then assembled and quantified using the StringTie software (version 2.1.5) (79). Differential gene expression analysis was performed using the *R* package DESeq2 (80) and the search for transcripts corresponding to gene fusion events was performed using the STAR-Fusion software assessed in (81). SAMtools (82) was used for transcript sorting, generation of files in pileup format and interconversion between BAM and SAM file formats.

### Preparation of scRNA-seq libraries

Frozen cells were thawed in prewarmed complete media (IMDMM with 10% FBS) and washed twice in complete media. The cell pellet was resuspended and cells stained with DAPI (Thermo Scientific, Product #62248, 1:1000 dilution) to exclude dead cells during cell sorting, and samples were collected in complete media. Sorted cells were washed and suspended in cold PBS + 0.04% BSA at optimal density before loading the 10X chromium controller system (10X Genomics Inc., product code 120223). Cells were barcoded using the 10X GenCode Technology and transcriptomes loaded onto Gel Bead-in-Emulsions (GEMs) and RT reactions, cDNA amplification, fragmentation, end repair and A tailing was performed as per manual instruction to obtain final libraries containing the P5 and P7 priming sites used in IlluminaR sequencing. High sensitivity DNA chips and Agilent 2100 bioanalyzer (Agilent Technologies) were used for 5’ gene expression quality control and quantification, performed twice before sequencing. The 5’ gene expression library was sequenced on NovaSeq 6000 S1 with sequencing depth of approximately 300-500 million reads per sample.

### Processing and analysis of scRNA-seq libraries

Illumina sequencing output files were converted to fastq format, scRNA-seq reads aligned to the human genome assembly GRCh38 (release version 84), and count matrix of cell barcodes by genes generated using Cell Ranger version 3.0.2 (10X Genomics). Filtered count matrices generated by Cell Ranger were loaded directly into Seurat software version 3.2.2 (38) for initial quality control where transcripts for genes only detected in three or less cells and cells with 200 or less transcripts from unique genes, or greater than 15% transcripts derived from mitochondrial genes, were excluded from further analysis. At this stage we found a subset of cells, subsequently assigned to the MitoLow plasma cell cluster, could be identified as clear outliers with less 3% of transcripts derived from mitochondrial genes. We used Seurat to perform log-normalization (scale factor = 10,000), variable feature selection (number of features = 2000), principle component analysis (PCA) and unsupervised Louvain clustering (resolution of 0.5) with the top twenty principle components, resulting in fourteen initial cell clusters. Initial clusters were then parsimoniously separated into three, mutually-exclusive groups that together accounted for all 5610 cells, based on the expression of the cell markers *EPCAM* (epithelial cell marker), *CD45* (immune cell marker) or *FAP* (fibroblast cell marker) by at least 20% of all cells within a cluster. Cells pooled from *CD45* clusters then underwent a second round of feature selection, PCA and clustering (resolution of 1.0). To identify cell types, a custom function written in *R* was used to quantify the number of cells in each cluster expressing particular marker genes, and those where at least 20% of cells expressing *EPCAM, CD45, FAP, CD14, CD79A, SDC1, CD3D*, and/or *CD8A* were annotated accordingly. All differential expression analyses were conducted using the *FindMarkers* function in Seurat that implements Wilcoxon signed-rank testing, and a customized version single-cell gene set analysis was performed by adapting the Seurat function *AddModuleScore*, sequentially applying it to the Seurat object with each gene set supplied as a list of features. Differential module scores (gene set scores) between clusters were quantified and assigned an adjusted p-value using the *FindMarkers* function. To correlate expression profiles of immune cell clusters with those from (31), the Seurat function *AverageExpression* was used to calculate the average expression profile for each immune cluster based only on genes with transcripts detected in both data sets. Subsequently, pairwise correlations were calculated between average expression profiles of each pair of clusters from each study.

### Metabolic profiling and analysis

Metabolic profiling was performed in collaboration with Metabolon Inc. as described in (83). Normalized metabolite abundances for ccRCC tumor samples, adjacent normal tissue samples and the single RMC tumor sample were merged together into a single data set that was pre-processed to remove measurements corresponding to any metabolite that was not simultaneously observed in all samples. Abundances were then transformed to Z-scores by subtracting mean abundance for given metabolite and dividing by standard deviation prior to principle component analysis (PCA) using the *R* programming language. Data were separated into tumor and normal tissue subsets, Z-scores re-calculated for each group, and the Crawford-Howell test was applied to each metabolite individually as implemented in the *R* package *psycho* using the RMC tumor or paired adjacent normal medullary tissue as a test value for tumor or normal tissue group, respectively. Reported p-values were adjusted using the Bonferroni Correction and metabolites with significantly different Z-scores (defined by an adjusted p-value cutoff value of 0.05) identified as reported in the Results section.

## Data and code availability

Sequencing data will be released alongside the final published version of this article. Code for reproducing the analyses is available at the GitLab repository https://gitlab.com/davidtourigny/renal-medullary-carcinoma

## Acknowledgments

This work was supported in part by NIH grant number P30 CA008748 and US DOD grant number W81XWG-18-1-0318. E.R. is the recipient of a Cycle for Survival Equinox Innovation Award, Brown Performance Group and KCA Young Investigator Award. R.R.K. is supported (in part) by the Academy of Kidney Cancer Investigators of the CDMRP/DOD (KC200127).

## Supplementary Material

**Table S1**. Results of differential gene expression analysis for all cell clusters.

**Table S2**. Results of differential gene expression analysis for three sub-clusters of epithelial-like cells.

**Table S3**. Results of Crawford-Howell test for all metabolites from tumor samples.

**Table S4**. Results of Crawford-Howell test for all metabolites from normal tissue samples.

